# The plasma membrane as a competitive inhibitor and positive allosteric modulator of KRas4B signaling

**DOI:** 10.1101/809616

**Authors:** C. Neale, A.E. García

## Abstract

Mutant Ras proteins are important drivers of human cancers, yet no approved drugs act directly on this difficult target. Over the last decade, the idea has emerged that oncogenic signaling can be diminished by molecules that drive Ras into orientations in which effector binding interfaces are occluded by the cell membrane. To support this approach to drug discovery, we characterize the orientational preferences of membrane-bound K-Ras4B in 1.45 milliseconds aggregate time of atomistic molecular dynamics simulations. Individual simulations probe active or inactive states of Ras on membranes with or without anionic lipids. We find that the membrane orientation of Ras is relatively insensitive to its bound guanine nucleotide and activation state but depends strongly on interactions with anionic phosphatidylserine lipids. These lipids slow Ras’ translational and orientational diffusion and promote a discrete population in which small changes in orientation control Ras’ competence to bind multiple regulator and effector proteins. Our results suggest that compound-directed conversion of constitutively active mutant Ras into functionally inactive forms may be accessible via subtle perturbations of Ras’ orientational preferences at the membrane surface.

**Statement of Significance:** Mutations that lock Ras proteins in active states can undermine cellular decision making and drive cancer. Because there are no drugs to deactivate Ras, we use simulations to relate Ras’ three-dimensional orientation at the membrane surface to its signaling competence. Data shows that Ras reorientation is generally rapid, but can be trapped in one of three states by membrane adhesion of the globular signaling domain. One of these states is stabilized by negatively charged lipids and brings an effector binding interface toward the membrane surface, potentially obstructing protein-protein interactions required for propagation of the growth signal. Rare events drive a second type of membrane-based signaling obstruction that correlate with configurational changes in Ras’ globular domain, yielding a potential drug target.

## Introduction

Ras proteins are intracellular signal transducers (1) that undergo regulated transitions between active and inactive states based on the identity of a bound guanine nucleotide (2). Whereas guanosine diphosphate (GDP) stabilizes inactive conformations, guanosine triphosphate (GTP) promotes active configurations that can bind downstream effectors (3).

Healthful signaling through Ras depends on its balanced response to upstream regulatory signals. Activation of Ras is accomplished by guanine nucleotide exchange factor proteins (GEFs) that accelerate nucleotide release to facilitate subsequent GTP binding (4, 5). The reverse process of deactivation occurs via enzymatic cleavage of GTP’s terminal phosphate group, which is dramatically accelerated upon binding to GTPase activating proteins (GAPs) (6, 7). However, oncogenic mutations in Ras proteins reduce the rate of GAP-stimulated deactivation (6, 8), leading to abnormally large populations of active Ras. Because Ras acts within signaling cascades that control cellular growth and survival (9-14), these overactive mutant Ras proteins are common drivers of human cancers (15-18).

The activation state of Ras is defined in part by the arrangement of a ∼10 residue component of Ras’ effector-binding interface called switch 1 (3, 19-21). Crystal structures of Ras bound to nonhydrolyzable GTP analogs reveal that the active state involves close association of switch 1 with other parts of Ras’ globular G domain (22, 23), including interactions between switch 1 residue T35 and G domain-bound GTP/Mg^2+^ (2). When Ras is bound to GTP or an analog, ^31^P NMR experiments resolve two interconverting conformational ensembles called states 1 and 2 (24-27). These states likely represent inactive and active Ras, respectively, because state 1 is stabilized upon binding to the catalytic site of a GEF (28) and state 2 is stabilized by effector binding (24-28). Deactivation of Ras involves disordering of switch 1, which has been characterized for wild-type Ras bound to GDP (29) and an inactive T35S mutant bound to a GTP analog (30, 31). Inactive nucleotide-free Ras bound to the GEF SOS1 crystalizes with switch 1 splayed more than 1 nm away from the nucleotide binding site (32).

Ras signals at the plasma membrane, to which it is localized by a C-terminal extension called the highly-variable region or HVR (33, 34). This localization adds an additional layer of complexity to the instantaneous definition of activity for a given molecule of Ras. Specifically, it is possible for Ras to bind GTP, adopt active structures, and yet be oriented such that the membrane occludes interactions with other proteins and signaling is temporarily obstructed. There is substantial evidence that Ras proteins adopt particular sets of orientations at membrane surfaces (35-41). These orientational preferences are modulated by lipid composition (37) and the identity of the bound nucleotide (35, 42-45), with clear functional relevance (42, 45-47). Indeed, a small molecule has recently been shown to stabilize Ras orientations that are incompetent to bind Raf kinase, an important downstream effector (48). However, the molecular mechanisms by which Ras’ bound nucleotide and/or activation state influence membrane interactions remain elusive.

Here, we report atomistic molecular dynamics (MD) simulations of full-length, post-translationally processed K-Ras4B (hereafter Ras) tethered to a lipid bilayer by its farnesylated HVR. We use hundreds of 5 μs simulations totaling 1.45 ms to characterize the influence of nucleotide identity, activation state, and lipid composition on the dynamic orientation of Ras and its interactions with membrane surfaces. Multiple simulations are needed to obtain statistically significant orientational distributions. We use the resulting orientational maps to assess the activity of the cell membrane as a competitive inhibitor or a positive allosteric modulator that reversibly blocks protein complexation or preconditions Ras for binding to other signaling proteins, respectively.

This research is part of a collaboration between the U.S. Department of Energy and National Cancer Institute to accelerate the development of novel diagnostics and targeted therapies for Ras-driven cancer initiation and growth.

## Methods

Simulation systems consist of a hydrated lipid bilayer with a single molecule of full length K-Ras4B tethered to one of the bilayer leaflets via its farnesylated HVR. The nucleotide binding pocket contains a magnesium ion and either GDP or GTP, with switch 1 initially in inactive or active configurations, respectively. N-terminal residue M1 has a backbone NH_3_^+^. C-terminal residue C185 is backbone-methylated and side chain-farnesylated. Lipid bilayers are composed of either pure 1-palmitoyl-2-oleoyl-*sn*-glycero-3-phosphocholine (POPC) or a 7:3 ratio of POPC and 1-palmitoyl-2-oleoyl-*sn*-glycero-3-phospho-L-serine (POPS). Simulation systems have 200 lipids per bilayer leaflet, 150 mM excess KCl, and ∼115 water molecules per lipid, consistent with a fully hydrated lipid bilayer (49). The G domain and HVR comprise residues M1-H166 and K167-C185, respectively.

### Simulation Parameters

Simulations are conducted with mixed-precision (SPFP (50)) AMBER 16 software (51). Protein and lipids are modeled by the CHARMM36 force field with CMAP (52, 53). All histidines are the neutral tautomer with a proton on the epsilon nitrogen. C-terminal methylation parameters are from the CHARMM CT1 patch. C-terminal farnesyl parameters are from Neale and García (54). Guanine nucleotide parameters are based on guanosine monophosphate and pyrophosphate parameters developed for use with CHARMM36 nucleic acids (55). Magnesium parameters are those introduced for use with CHARMM27 nucleic acids (56). The water model is TIP3P (57) with CHARMM modifications (58). AMBER formatted topologies are obtained with the gromber tool of ParmEd from AmberTools 16 after initial topology construction with GROMACS 5.1.2 (59). Water molecules are rigidified with SETTLE (60) and other covalent bond lengths involving hydrogen are constrained with SHAKE (61) (tolerance=10^−6^ nm). Lennard-Jones (LJ) interactions are evaluated using an atom-based cutoff with forces switched smoothly to zero between 1.0 and 1.2 nm. This is the recommended LJ cutoff for the CHARMM36 protein force field (52, 62). Note that the CHARMM36 lipid force field was parameterized with LJ switching between 0.8 and 1.2 nm (53). Coulomb interactions are calculated using the smooth particle-mesh Ewald method (63, 64) with Fourier grid spacing of 0.08 to 0.10 nm and fourth order interpolation. Simulation in the *NpT* ensemble is achieved by semi-isotropic coupling to Monte Carlo barostats (51) at 1.01325 bar with compressibilities of 4.5×10^−5^ bar^−1^; temperature-coupling is achieved using velocity Langevin dynamics (65) at 310 K with a coupling constant of 1 ps. The integration time step is 4 fs, which is enabled by hydrogen mass repartitioning (66). Non-bonded neighbor-lists are built to 1.4 nm and updated heuristically (51).

### Simulation set

We conduct a total of 290 5-μs simulations. As outlined in Table 1, 200 of these simulations contain inactive, GDP-bound Ras, of which 116 use a POPC bilayer and 84 use a mixed 7:3 POPC:POPS bilayer. The other 90 simulations contain initially active, GTP-bound Ras on a mixed 7:3 POPC:POPS bilayer. We do not simulate active, GTP-bound Ras with pure POPC bilayers.

**Table 1:**
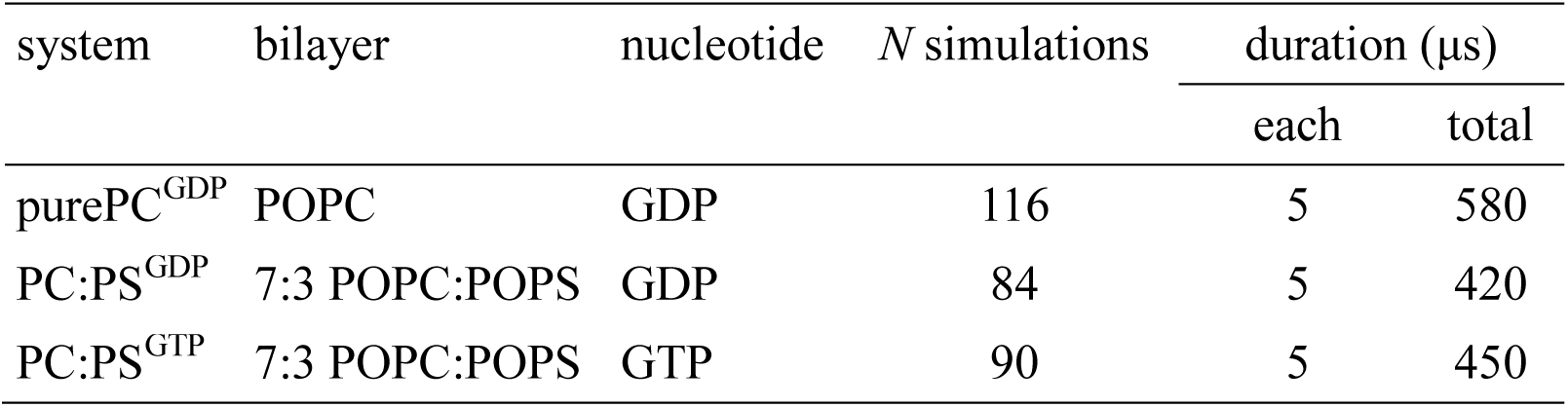
Simulation systems.

Systems with 195,000 ± 5,000 atoms are constructed by adding G domains to an equilibrated ensemble of HVR configurations drawn from our previous work (54). Simulations of GDP- and GTP-bound Ras are based on 4OBE (67) and 5USJ (68) structures, respectively. Details of system construction are provided in SI Methods.

### Order parameters

G domain orientation is defined by tilt and rotation angles (69) with respect to a reference orientation in which helix 5 (α5; V152-K165) from chain A of PDB 4OBE (67) is aligned with the global bilayer normal (Cartesian **z** axis). This reference orientation is compared to each sampled orientation to compute its tilt away from **z** and the azimuthal angle of that tilt (i.e., the rotation angle) using the rotation matrix that minimizes the root mean squared deviation between C_α_ atoms of G domain residues T2-N26, Y40-L56, and D69-H166. Tilting toward the center of mass of the C_α_ atoms of α2 (S65-T74) is defined as rotation=0°. Viewing the reference orientation from N-terminal to C-terminal end of α5 (i.e., toward the membrane plane), positive and negative rotations involve tilting toward G domain regions counter-clockwise and clockwise from α2, respectively.

### Analysis

Time averaged data are computed from the 2-5 μs trajectory segment (the last 3 μs) in each simulation. Orientational “fold enrichment” is obtained by dividing the probability at which each histogram bin is sampled by the per-bin probability expected from a homogeneous distribution over 180° of tilt and 360° of rotation with bin widths of 3° and 6°, respectively. This approach corrects for variable histogram bin size as a function of tilt angle without introducing a less intuitive order parameter such as cos(*tilt*). Orientations of published structures are computed from raw coordinates, except models from PDBs 6CCH, 6CC9, and 6CCX (48), which are first aligned to the nanodisk belt proteins in model 1 from PDB 2MSC (44) so as to position the bilayer normal along the Cartesian **z** axis. Rotation angles for 2MSC are shifted by −12° to account for the α2 offset in model base PDB 4LPK (70) chain B vs. our reference structure (4OBE (67) chain A). Molar entropies of Ras G domain orientation are computed as *S*° = −*k*_B_×∑*P*_*tilt,rot*_ × ln (*P*_*tilt,rot*_) for Boltzmann constant *k*_B_ and non-zero orientational probabilities *P*_*tilt,rot*_. To reduce noise in projections of orthogonal data onto maps of tilt and rotation, average values are only displayed for orientational bins that are sampled at least 5 times in both simulation subsets A and B (except the quantification of G domain capability to bind regulators and effectors, which shows all data). Our measure of membrane obstruction, *N*_clash_, is averaged over all trajectory frames from 2-5 μs in all 290 simulations after reorienting crystal structures to minimize the mean squared displacement of C_α_ atoms in G domain residues T2-N26, Y40-L56, and D69-H166 between simulated K-Ras4B and crystallographic H-Ras. Note that K-Ras and H-Ras G domains share a common fold (23) and have a gap-less sequence alignment with matching identity for 156/166 residues (71). Secondary structure is computed with DSSP version 2.0.4 (72).

Lateral diffusion coefficients are computed by fitting average intra-simulation Cartesian **x**,**y** plane mean squared displacements of the centers of mass of lipids or the Ras G domain to linear functions over Δ*t*=1-2 μs and dividing the fitted slope by 4. Simulation trajectories are processed to remove **x**,**y** motion of the center of mass of lipids in the bilayer leaflet under consideration prior to evaluating molecular diffusion (73). Diffusion coefficients computed from these simulations are not corrected for finite size effects stemming from hydrodynamic interactions (74-77). Since all systems have similar dimensions, we compare unmodified diffusion coefficients. Orientational displacement, *θ*, is the angle formed between two unit vectors in a three dimensional polar coordinate system of (*r*=1, tilt, rotation); i.e., *θ*_*i,j*_=cos^-1^(***V***_*i*_•***V***_*j*_) for vectors ***V***=[sin(*tilt*)×cos(*rot*),sin(*tilt*)×sin(*rot*),cos(*tilt*)] corresponding to orientations *i* and *j*. Survival probabilities, *S*(Δ*t*), are fit to 2^-Δ*t*/*τ*^, and the mean half life is reported as *τ*.

To estimate precision, simulations are divided into two sets and average values are computed for each set independently. Specifically, for *N* simulations of a given system composition, sets A and B comprise 2-5 μs trajectory segments from simulations [1,*N*/2] and [*N*/2+1,*N*], respectively. The standard deviation of the mean is calculated as 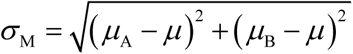, for subset mean values *μ*_A_ and *μ*_B_, and overall mean value *μ*. Block-averaging (78) represents ensemble average values from time *t* to *t* + Δ*t*.

## Results and Discussion

To study the influence of membrane composition on protein behavior, we simulate GDP-bound Ras with lipid bilayers composed of either POPC (purePC^GDP^ systems) or a 7:3 mixture of POPC:POPS (PC:PS^GDP^ systems). Nucleotide-dependent behavior is probed via comparison to simulations of GTP-bound Ras (PC:PS^GTP^ systems). A total simulation time of 1.45 ms is achieved in 290 5-μs simulations (Table 1). Ensemble averaged time series of Ras G domain center of mass distance from the bilayer and the number and type of G domain-lipid contacts lead us to discard the first two microseconds per simulation, after which ensembles of this size show no statistically significant drift in these metrics with increasing time (Fig. 1 and Table S1).

**Figure 1:**
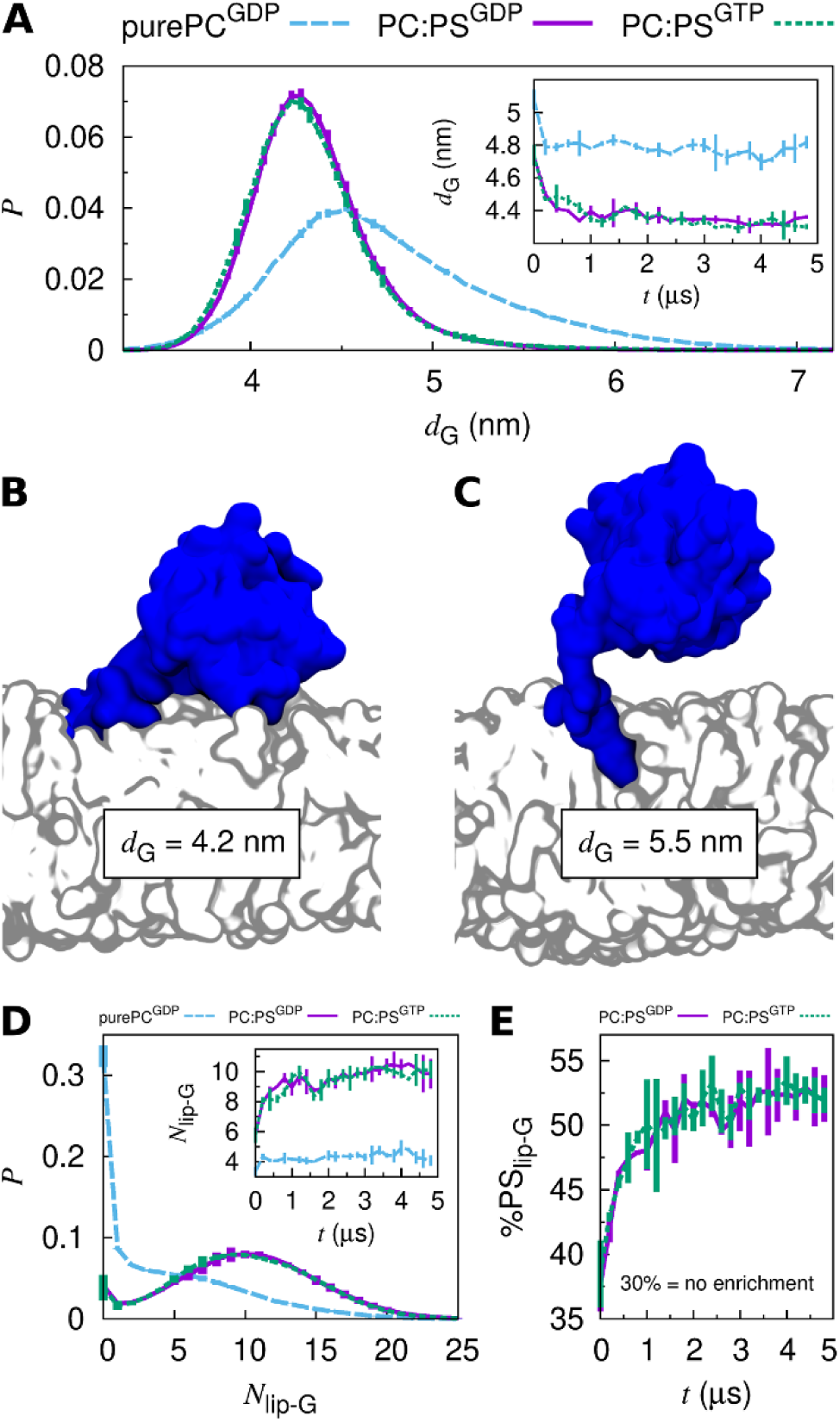
Ras G domain-membrane interactions. (A) Histograms of the distance between the centers of mass of the G domain and the bilayer along its normal, *d*_G_. Curves connect average values and bars show standard deviation of the mean. Inset: ensemble- and block-averaged time series of *d*_G_. (B and C) Representative configurations for (B) *d*_G_=4.2 nm and (C) *d*_G_=5.5 nm. (D) Histograms of the number of lipids contacting the G domain, *N*_lip-G_ (0.6 nm heavy atom cutoff). Inset: ensemble- and block-averaged time series of *N*_lip-G_. (E) Ensemble- and block-averaged time series of the percentage of all lipids contacting the G domain that are PS lipids, %PS_lip-G_.

### Membrane Adhesion and Lateral Diffusion

Simulations of full-length Ras are largely consistent with our previous HVR-only simulations (54) in which anionic PS lipids attract the HVR, and therefore the G domain, toward the membrane (Figs. 1A-C and S1). In the context of full-length Ras, we find that the identity of the bound nucleotide does not significantly affect Ras’ displacement from the bilayer surface (Fig. 1A). In contrast, PS lipids reduce the likelihood of G domain disengagement from the lipid bilayer by a factor of 8 ± 1 (Fig. 1D) and are locally enriched by a factor of 1.7 ± 0.1 (Fig. 1E). Concurrently, PS lipids slow the lateral diffusion of Ras on the membrane surface by a factor of 2.2 ± 0.7 (Table 2).

**Table 2:**
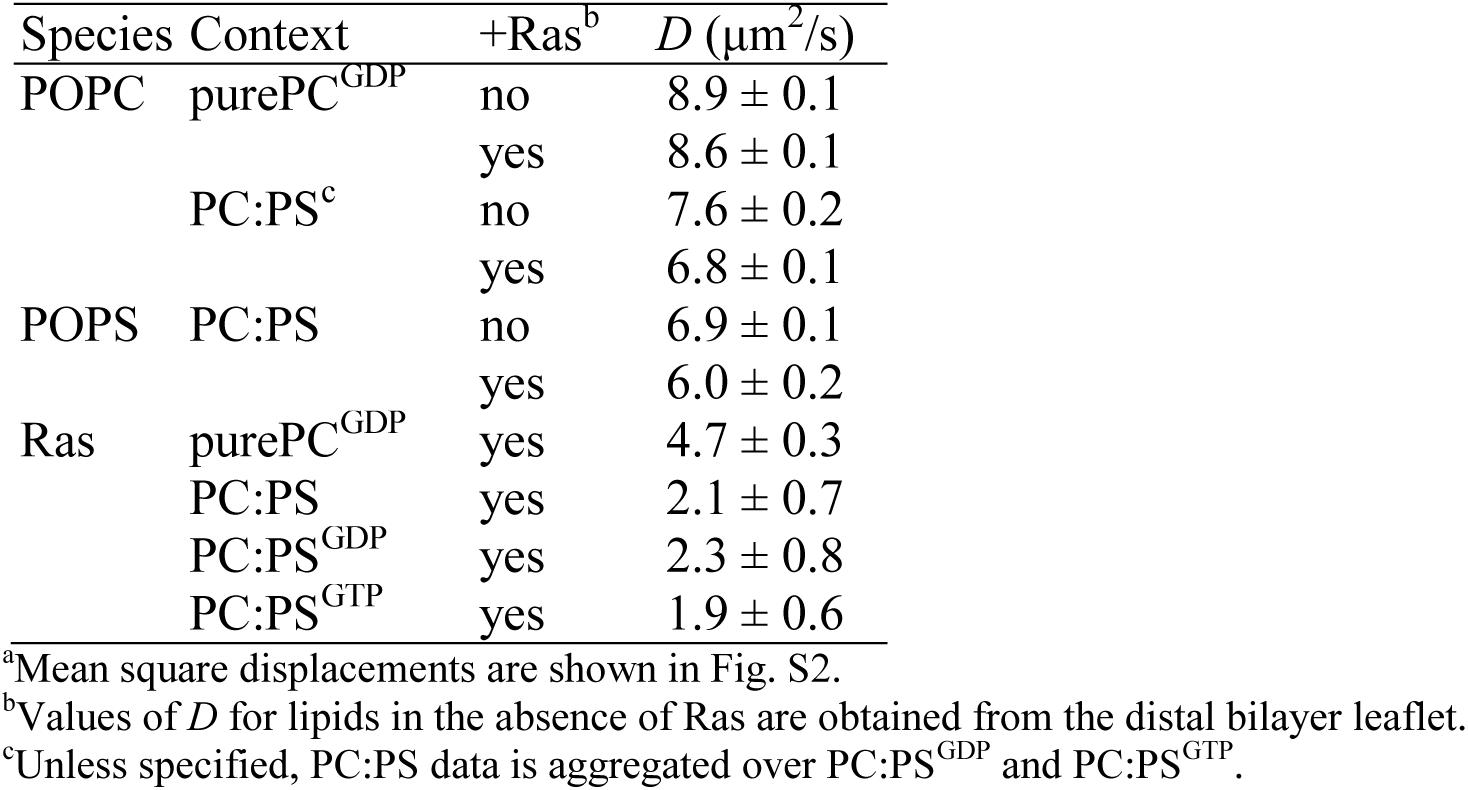
Lateral diffusion constants^a^, *D*.

The lateral diffusion constant we compute for Ras on 30% POPS bilayers (2.1 ± 0.7 μm^2^/s; Table 2) is slower than experimental rates from fluorescence correlation spectroscopy (∼4.5 μm^2^/s) and single particle tracking (∼3.5 μm^2^/s) in supported bilayers containing 20% 1,2-dioleoyl-*sn*-glycero-3-phospho-L-serine (DOPS) (79). This discrepancy may relate to our larger concentration of PS lipids, the minor population of slower diffusing (∼1 μm^2^/s) monomeric Ras identified by single particle tracking (79), or finite size effects in our simulations (see Methods). Nevertheless, simulation (Table 2) and experiment (79) agree that diffusion rates are nucleotide-independent.

### Orientation with Respect to the Bilayer Surface

We characterize the disposition of Ras’ G domain with respect to the bilayer surface using tilt and rotation angles (69). Briefly, the tilt angle describes the deflection of helix 5 (α5) from the bilayer normal, and the rotation angle describes the direction of the tilt (see Methods). Schematics identifying the regions of Ras that are brought into membrane contact upon tilting with different rotations are shown in Figs. 2A and 2B.

**Figure 2:**
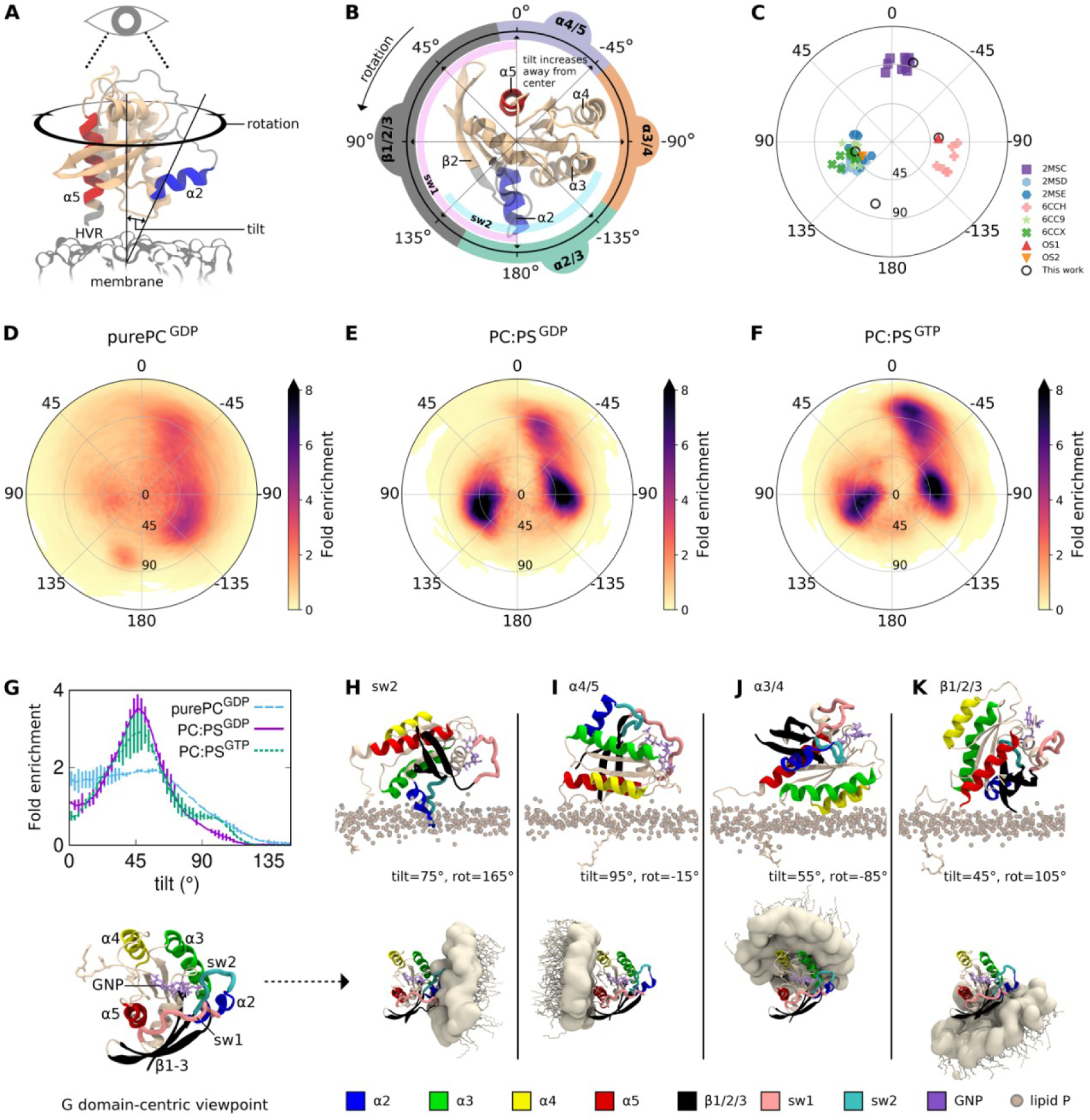
Orientations of the Ras G domain. (A) Reference orientation. (B) Rotation angles and corresponding elements of Ras brought into membrane contact upon tilting are inscribed around the reference orientation, viewed toward the bilayer along its normal. (C) Published orientations of membrane-bound K-Ras4B: PDBs 2MSC, 2MSD, and 2MSE are the “exposed”, “occluded”, and “semi-occluded” orientations from Mazhab-Jafari et al. (2MSE includes a bound Raf domain) (44); PDBs 6CCH, 6CC9, and 6CCX are the “E3”, “O1”, and “O2” orientations from Fang et al. (48); “OS1” and “OS2” are from Prakash et al. (36). (D-F) Heat maps of the fold enrichment over random for tilt and rotation angles sampled in this work. Statistical error is quantified in Fig. S3 and Table S2. Orientational definitions by alternative order parameters (36, 80) are provided in Figs. S4 and S5. (G) Fold enrichment of tilt angle. (H-K) Representative orientations that bring (H) switch 2, (I) α4/5, (J) α3/4, and (K) β1/2/3 into membrane contact, shown (above) with respect to the membrane plane and (below) in a G domain-centric frame of reference.

G domain orientations sampled in our simulations are shown in Figs. 2D-F. By comparing simulations with and without 30% PS lipids, we are able to characterize the lipid dependence of Ras orientation. Pure PC bilayers tend to engage with Ras’ helical α4/α5 or α3/α4 interfaces, and to a lesser extent with beta strands 1-3 and α2 (β1/2/3 for brevity) or switch 2 (Fig. 2D), the later plunging into the bilayer upon 75° tilting with a rotation angle near 165° (Fig. 2H). The addition of PS lipids further reduces the G domain’s orientational molar entropy, *S*°, which is 62.6 ± 0.1 J/mol/K for purePC^GDP^ systems vs. 58.1 ± 0.1 J/mol/K for PC:PS systems (computed from raw probabilities underlying Figs. 2D-F). This orientational restriction is consistent with the increase in G domain-lipid interactions upon addition of PS lipids (Fig. 1). In addition to favoring orientations in which Ras’ α4/α5 or α3/α4 interfaces interact with the membrane (Figs. 2E, F, I, J), these anionic lipids also direct Ras toward rotation angles near 105°, where tilting drives α2 and β1/2/3 toward the membrane (Figs. 2E, F, K). Although Ras tilts more extensively toward α4/α5 when bound to GTP compared to GDP (Figs. 2E, F, G), nucleotide-dependent differences in Ras orientation are of the same magnitude as statistical sampling errors (Fig. S3 and Table S2). Therefore, if Ras’ disposition does depend on the identity of the bound nucleotide, it cannot be determined from this extensive set of simulations.

Paramagnetic NMR in nanodisks with 20% PS lipids reveals the same three main states in which K-Ras4B tilting brings either α4/α5, α3/α4, or β1/2/3 into membrane contact (Fig. 2C) (44, 48). These states are also captured in published MD simulations (36, 41, 80), although the α4/α5 and α3/α4 states are not well resolved by alternative order parameters (Fig. S5). Together, these experimental and theoretical results indicate that Ras’ G domain favors distinct orientations at the membrane surface and that PS lipids modulates orientational preferences in a manner that may be druggable (48).

### Interactions with Lipids

All four population basins identified above involve G domain interaction with a greater than average number of lipids (Figs. 3A, B). In the PC:PS context, α4/5 orientations show close interactions between lipids and α4 residues K128 and R135 and, to a lesser extent, α5 residue R161 (Fig. 3C). α3/4 orientations also involve close lipid association with α4 residues K128 and R135, in addition to α3 residues R97, K101, and R102 (Fig. 3D). Finally, β1/2/3 orientations have close associations for β1 residues M1, E3, and K5, β2 residue R41, and α2 residue R73 (Fig. 3E). Lipid contacts observed in the α3/4 and β1/2/3 states are consistent with those previously identified via simulation (36). In the pure PC context, orientations with switch 2 membrane insertion show tight lipid interactions for switch 2 residues Y64, S65, R68, Y71, and R73 with less frequent interactions for α3 residue R102 (Fig. 3F).

**Figure 3:**
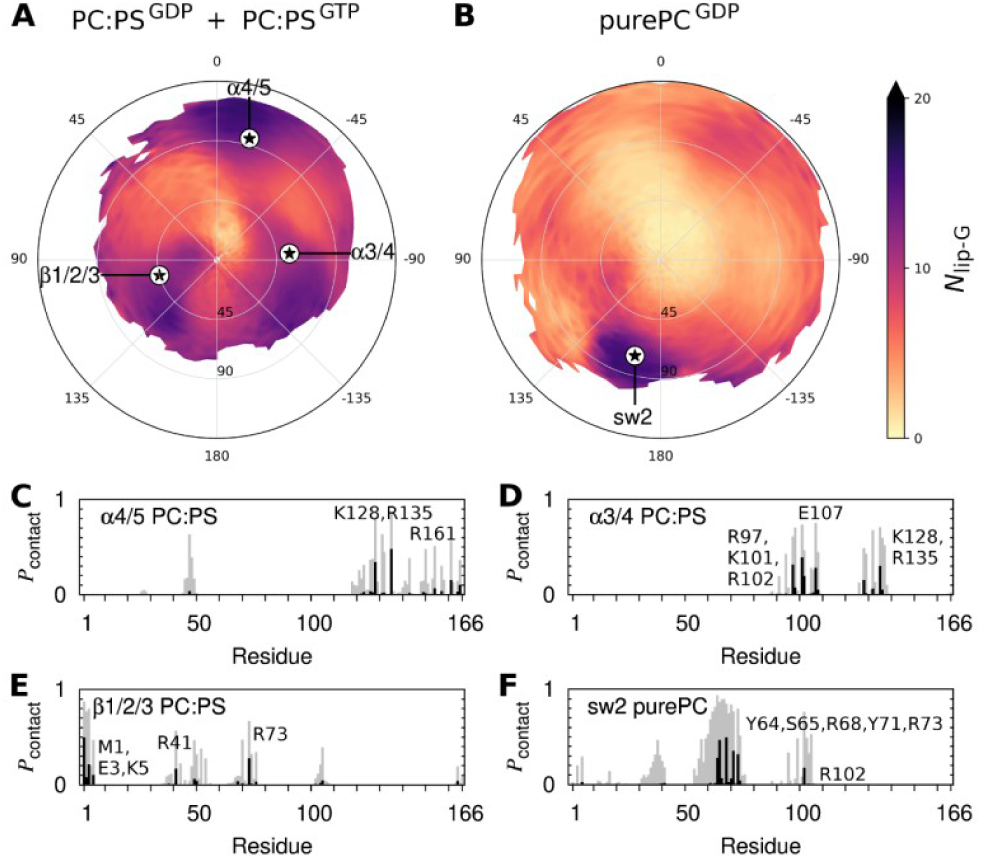
Orientational dependence of Ras G domain-membrane interactions. (A and B) Time and ensemble averaged values of the number of lipids contacting the G domain, *N*_lip-G_, for G domain orientations in simulations with (A) PC:PS and (B) PC lipids. Stars denote orientations shown in Figs. 2H-K, which are representative of the major populations evident in Figs. 2D-F. (C-F) Histograms showing the probability at which each G domain residue contacts a lipid in each of four different Ras orientations. Contacts are shown for heavy atoms closer than (gray) 0.6 nm and (black) 0.28 nm. Simulation snapshots are considered only when Ras’ tilt and rotation angles are each within 5° of the noted population peaks.

The imperfect alignment of highly populated orientations (Figs. 2D-F) with those that maximize the number of lipids contacting the G domain (Figs. 3A, B) is likely a geometrical consequence of HVR attachment to the C-terminal end of α5. By definition, this end of α5 points toward the membrane at tilt=0° and tilting correlates with membrane separation of C-terminal G domain residue H166, except when Ras tilts toward α4/5 (Fig. S6). Therefore, favored orientations of the G domain appear to be primarily driven by direct interactions with the membrane but are subject to constraints on tilting enforced by the HVR’s preference to associate with anionic lipids. Additional factors such as HVR-lipid contacts (Fig. S7C) and HVR-G domain contacts (Fig. S7F) exhibit weaker correlations with highly populated orientations and may play minor roles in regulating G domain disposition.

The closer association of the G domain with the membrane in major population basins is also resolved by center of mass separation (Fig. S7A). Nevertheless, in the presence of PS lipids, the average minimum G domain-membrane separation is largely constant, suggesting that the G domain can remain in membrane contact as it rolls between α4/α5 and α3/α4, and between α3/α4 and β1/2/3 (Fig. S7B). In contrast, direct transitions between α4/α5 and β1/2/3 are more likely to involve transient disengagement of the G domain from the membrane (Fig. S7B). This feature may result from unfavorable interactions between lipids and the anionic β1/ β2 loop (which contains D47 and E49), it’s convex shape (Fig. 2B), or strong dependence of α4/α5 membrane adhesion on α4 residues K128 and R135 (Fig. 3C), which must release the membrane during a rolling motion toward β1/2/3.

### Contact with the HVR

In contrast to the above mentioned relationships between orientational probability and G domain-membrane interactions, there are no obvious correlations with the number of lipids contacting the HVR (Fig. S7C) or anionic proportion among lipids in contact with the G domain (Fig. S7D) or the HVR (Fig. S7E). Although the number of HVR residues contacting the G domain is elevated in orientations near the α4/5 and α3/4 states (Fig. S7F), G domain-HVR interactions are reasonably well predicted by models in which contact probabilities fall off linearly with increasing HVR residue number and, within the G domain, based on rigid-body distance from the HVR attachment point at H166 (Fig. S8). Therefore, our simulations of farnesylated, membrane bound Ras are not consistent with a unique, high affinity HVR binding site on the G domain, as has been proposed for autoinhibition of GDP-bound K-Ras4B in the absence of a membrane (81, 82).

### Length of G domain Helix 5

Circular dichroism spectra of K-Ras4B HVR peptides associated with anionic membranes are consistent with structural disorder (83), as are our previous simulations (54). Nevertheless, some crystal structures resolve α5 helical extension to HVR residue K172 (84) or D173 (85) and there is extensive helical content in the HVR of full-length K-Ras4B in complex with PDEδ (86). It is therefore an open question whether the G domain nucleates helical extension into the HVR, especially since nearly all crystal structures of K-Ras4B are based on truncated sequences containing only the G domain. Irrespective of nucleotide identity or the presence PS lipids, our simulations reveal that α5 loses 50% of its helicity at residue K167, with extension to K169 or S171 in 16 ± 2% and 5 ± 1% of the population, respectively (Fig. 4A). Although there is no simple relation between α5 length and G domain orientation in simulations with PC:PS lipids (Fig. 4B), switch 2 embedding in pure PC bilayers appears to correlate with C-terminal extension of α5 (Fig. 4C-E). Nevertheless, only 3 of 116 purePC^GDP^ simulations exhibited switch 2 embedding for more than 1 μs and therefore this state has less statistical significance than others.

**Figure 4:**
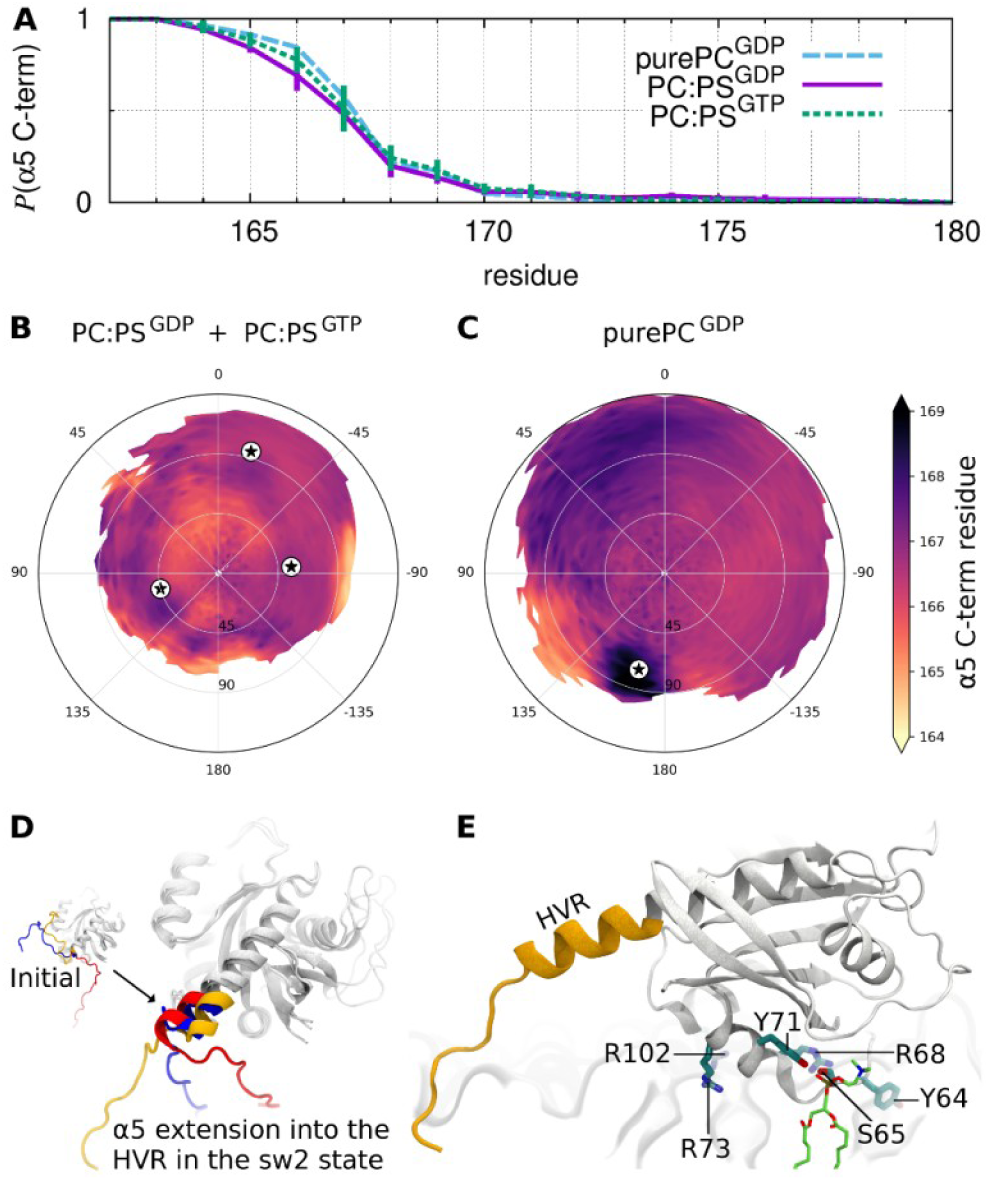
Length of the HVR-proximal G domain helix, α5. (A) Probability at which α5 extends to the noted residue. (B and C) Time- and ensemble-averaged values of the C-terminal residue in α5 as a function of G domain orientation. (D) Overlaid snapshots of three simulations in which α5 initially terminated at or near H166 and later adopted two additional helical turns into the HVR with concurrent membrane-embedding of switch 2. (E) Representative configuration with membrane-embedded switch 2, highlighting lipid interaction with S65 and Y71.

### Membrane Insertion of HVR residue M170

In our previous simulation study of isolated HVR peptides, we proposed a fly-casting mechanism by which the HVR’s poly-lysine stretch (K_175_KKKKK_180_) attaches to anionic lipids whereas HVR residue M170 seeks out membrane defects (54). Consistent with those results, the present simulations show that spontaneous membrane insertion of M170 is possible in the context of full-length K-Ras4B (Fig. 5). This interaction is more common in the presence of anionic PS lipids and is nearly exclusive to G domain orientations that are similar to the highly populated β1/2/3 state, excepting a reduction in the tilt angle (Figs. 5A, B). In our set of 174 PC:PS simulations, 12 exhibited spontaneous M170 insertion that lasted for more than 1 μs. Nevertheless, the presence of the G domain reduces the likelihood of M170 membrane insertion, which is ∼25% for HVR peptides (54) and only 4 ± 2% in the context of full-length Ras. This attenuation likely arises from spatial constraints given that, in the presence of PS lipids, the G domain forces this region of the HVR an average of ∼0.5 nm farther from the membrane surface, whereas the disposition of other regions of the HVR are relatively unaffected (Fig. S1).

**Figure 5:**
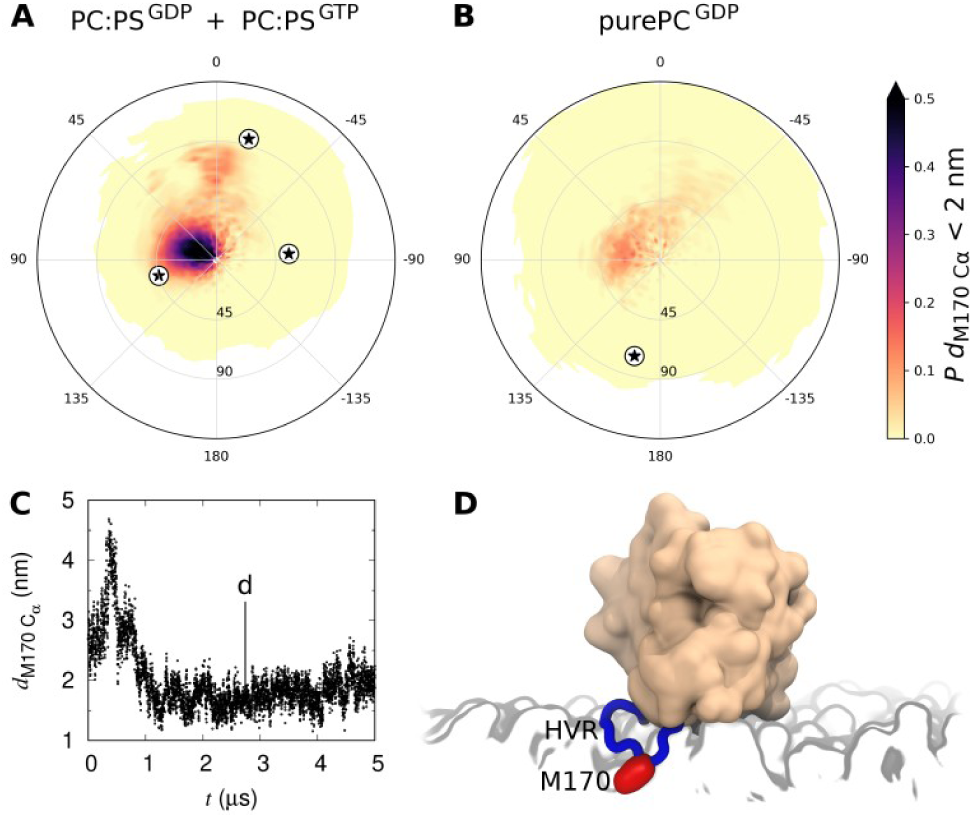
Membrane insertion of HVR residue M170. (A and B) Probability of the M170 C_α_ atom being closer than 2.0 nm to the bilayer center along its normal. (C) M170 C_α_ distance from the bilayer center in a representative PC:PS simulation in which M170 undergoes spontaneous membrane insertion. (D) Snapshot from the trajectory shown in part C.

### G Domain Reorientation

In addition to modulating orientational preferences and slowing lateral diffusion, anionic PS lipids slow the G domain’s orientational diffusion by an order of magnitude (Fig. 6A). As was the case for lateral diffusion, rates of orientational diffusion do not depend on nucleotide identity (Fig. 6A). Comparison to average orientational displacement between two randomly selected snapshots from the same simulation set shows that this effect is kinetic (Fig. 6B) and does not simply arise from the time and ensemble averaged orientational limitations imposed by PS lipids (Fig. 2). Importantly, time-dependent orientational displacement is smaller and does not depend on lipid composition when evaluation is restricted to trajectory segments in which the G domain maintains at least one lipid contact (heavy atom distance <0.28 nm; Fig. 6C). In fact, average orientational displacements between times *t* and *t*+0.5 μs depend linearly on the largest G domain-lipid separation distance encountered (Fig. 6D). Transient excursions of the G domain away from the bilayer surface are more common in the absence of PS lipids (Figs. 1A, 1D, and S9), except when switch 2 is embedded in the bilayer (Fig. 6E). Similarly, the α4/α5, α3/α4, and β1/2/3 states change orientation more slowly than intervening orientations in the presence of PS lipids (Figs. 6F, G). Together, these results suggest that a major component of G domain reorientation occurs during its transient disengagement from the membrane, after which reassociation is enforced by the still-tethered HVR.

**Figure 6:**
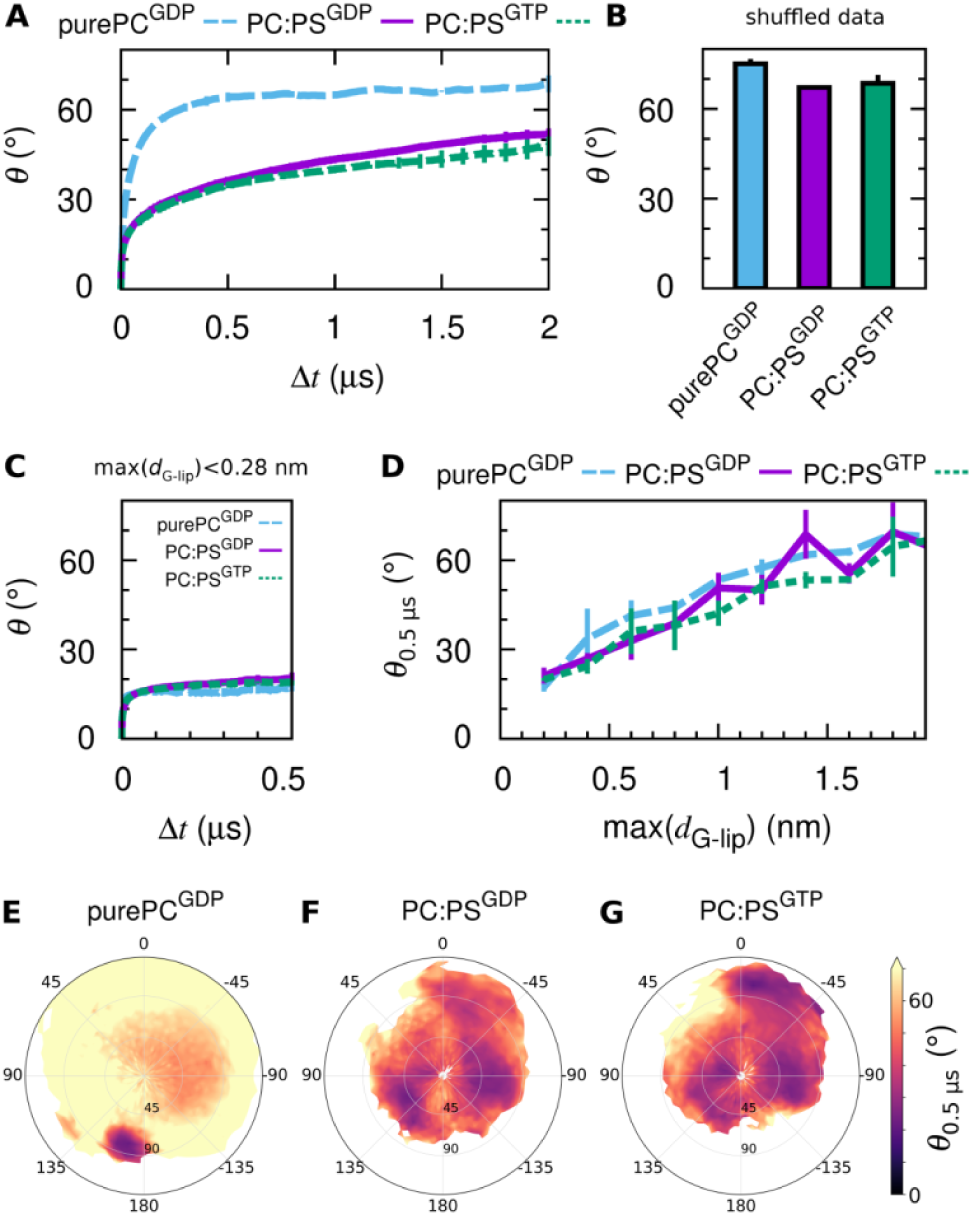
Orientational diffusion of the Ras G domain. (A) Average orientational displacement, *θ*, between configurations at times *t* and *t*+Δ*t*. (B) Average value of *θ* from time- and simulation-shuffled data. (C) *θ* vs. Δ*t* for trajectory segments in which the G domain maintains lipid contact (minimum G domain-lipid separation, *d*_G-lip_<0.28 nm). (D) Average values of *θ* at Δ*t*=0.5 μs, *θ*_0.5μs_, as a function of the maximum value of *d*_G-lip_ in that time span. Analogous plots for different time spans are shown in Fig. S10. (E-G) *θ*_0.5μs_ as a function of G domain orientation at time *t*.

### Active State Decay

Our simulations of GTP- and GDP-bound Ras are designed to probe the influence of activation state on membrane interaction under the assumption that the presence of GTP is sufficient to maintain active Ras configurations. However, out of 90 PC:PS^GTP^ simulations, 21 incur spontaneous detachment of switch 1 residue T35 from GTP/Mg^2+^ (Figs. 7A, B). This detachment does not conspicuously depend on Ras orientation (Fig. 7C). Survival probabilities indicate a 11 ± 2 μs half life of the T35-Mg^2+^ interaction (Fig. 7D), which is much shorter than the 90 μs half life of active-like configurations of switch 1 at 310 K calculated based on ^31^P NMR for H-Ras complexed with the GTP analog Gpp(NH)p (24). Moreover, conformational active-state half lives are an order of magnitude larger for GTP-compared to Gpp(NH)p-bound Ras (27). Therefore, our simulations underestimate the stability of Ras’ active configurational ensemble by up to two orders of magnitude. Nevertheless, orientations sampled in these 21 simulations are similar to those in 69 PC:PS^GTP^ simulations in which T35 remains coordinated by GTP/Mg^2+^, supporting our conclusion that the membrane orientation of K-Ras4B does not substantially depend on the identity of the bound nucleotide or the arrangement of switch 1 (Figs. 7E, F).

**Figure 7:**
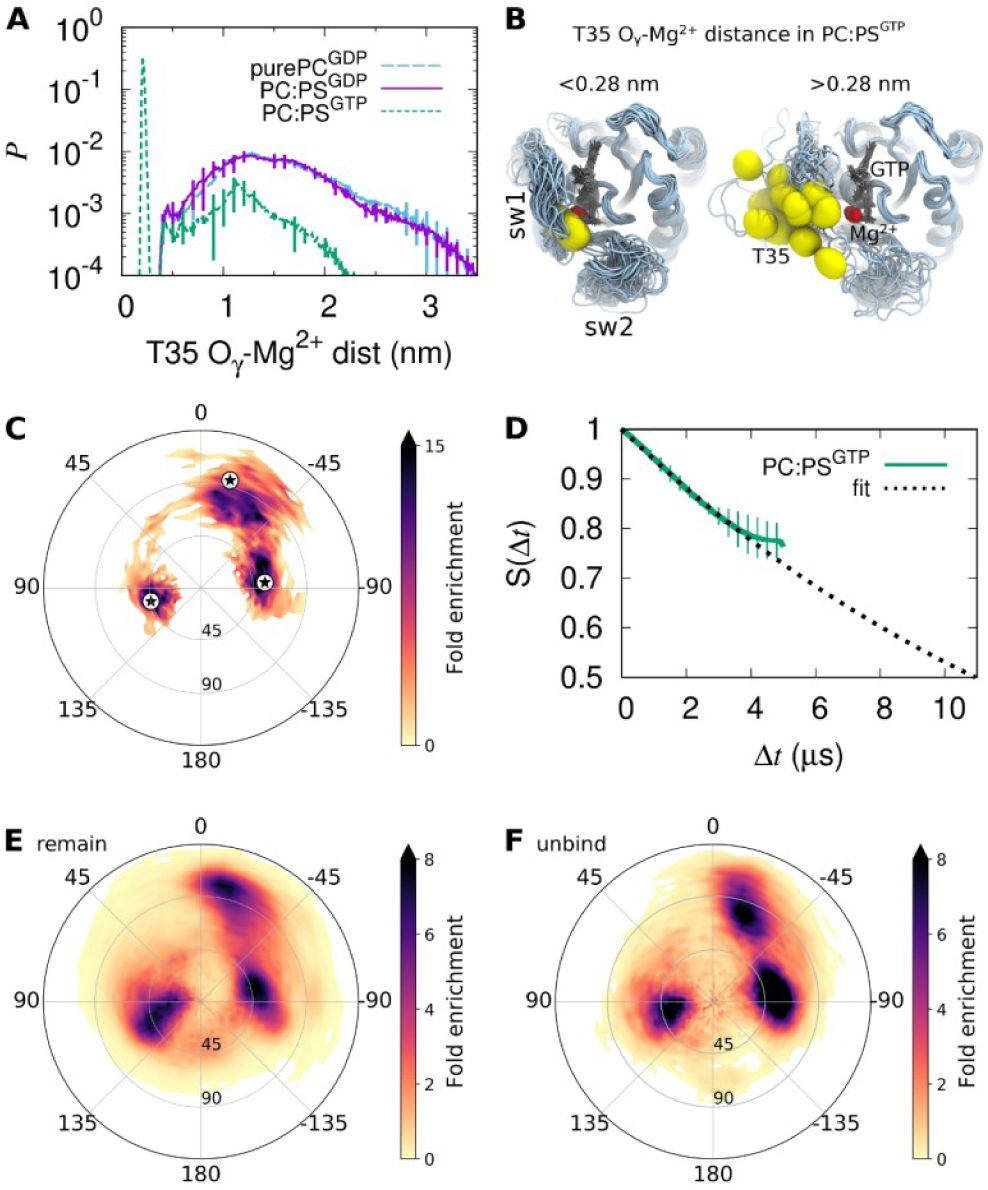
Detachment of switch 1 in 21 of 90 PC:PS^GTP^ simulations. (A) Probability histograms of the distance between T35 side chain oxygen atom O_γ_ and the GTP-coordinated Mg^2+^ ion. On the 5 μs/simulation timescale, distances <0.28 nm are maintained in 83±4% of the sampling for PC:PS^GTP^ systems. Approaches closer than 0.28 nm are never observed in GDP systems. (B) Overlay of final configurations in (left) 69 simulations with stable T35-Mg^2+^ interaction and (right) 21 simulations with T35 unbinding. (C) Orientations sampled within ±100 ns of T35-Mg^2+^ unbinding. (D) Survival of the T35-Mg^2+^ interaction fit to an exponential decay (0.28 nm interaction cutoff). (E and F) Heat maps of the fold enrichment over random for tilt and rotation angles sampled in PC:PS^GTP^ simulations in which the T35 side chain oxygen (E) did not, or (F) did dissociate from the GTP-coordinated Mg^2+^ ion.

### Binding to Regulators and Effectors

Interest in Ras’ orientation is largely due to its potential impact on the binding of regulators and effectors. In this context, the lipid bilayer may act as a competitive inhibitor that binds reversibly to Ras interfaces, thereby occluding them. To quantify the orientational dependence of Ras’ competence to bind other proteins, we align crystal complexes on Ras G domain configurations from our simulations and assess the spatial overlap between the crystallographic binding partner and the simulated lipid bilayer (Fig. 8). Clearly, none of these binding partners can engage Ras when switch 2 is embedded in the bilayer. Moreover, P120GAP binding is inconsistent with the majority of α3/α4 orientations and the allosteric Ras bound to SOS1 cannot adopt most α4/α5 orientations. Despite the appearance that most proteins can bind α3/α4- and α4/α5-oriented Ras, these binding partners are typically much larger than their co-crystalized domain and therefore orientations that are predicted to be inaccessible for binding via this approach generally represent subsets of the true orientational restrictions.

**Figure 8:**
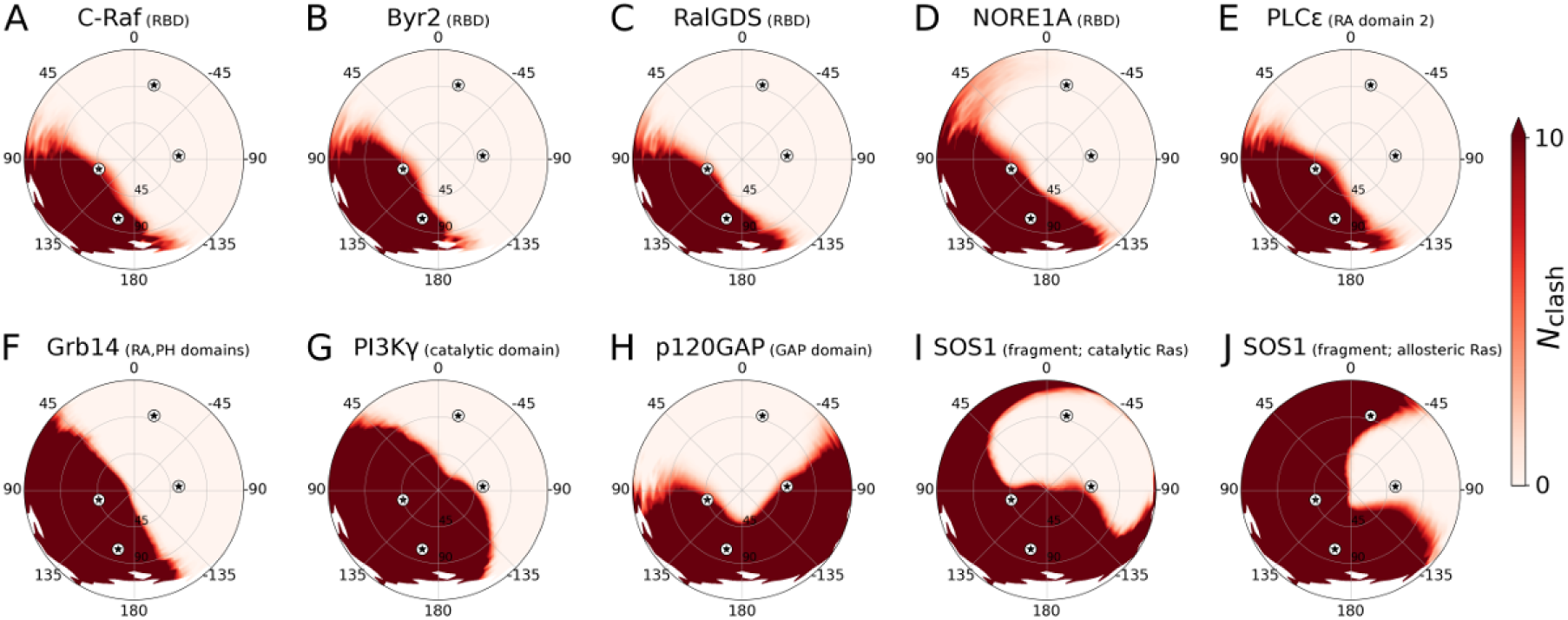
Influence of orientation on the G domain’s ability to bind regulators and effectors. To evaluate the spatial accessibility of Ras’ binding sites for regulators and effectors, crystal structures of H-Ras in complex with another protein are aligned on K-Ras4B G domains from our simulations. Membrane obstruction of this binding is then computed as the number of C_α_ atoms in the binding partner that are closer than 1.8 nm to the bilayer center, *N*_clash_, whose average value is plotted as a function of Ras G domain tilt and rotation. PDB structures used in this evaluation are (A) The Ras-binding domain (RBD) of C-Raf (4G0N) (3), (B) the Byr2 RBD (1K8R) (19), (C) the RalGDS RBD (1LFD chains A and B) (20), (D) the NORE1A RBD (3DDC) (21), (E) the PLCε Ras-associating (RA) domain 2 (2C5L chains A and C) (87), (F) the Grb14 RA and pleckstrin-homology (PH) domains (4K81 chains A and B) (88), (G) the PI3Kγ catalytic domain (1HE8) (89), (H) the p120GAP GAP domain (1WQ1) (8), and (I and J) a SOS1 fragment (residues 566-1046), containing two Ras binding sites (1NVV) (32). Part (I) shows the ability of SOS1 to bind Ras via its active site and part (J) shows Ras binding to the allosteric site on SOS1. Note that actual values of *N*_clash_ may be larger in cases where the binding protein is only partially represented by the crystal structure. As more details of these co-complexes are resolved, they can be assessed for possible further restriction of accessible binding orientations.

In addition to acting as a competitive inhibitor, the membrane may also act as a positive allosteric modulator by pre-orienting Ras for protein engagement. For example, Mazhab-Jafari et al. have constructed NMR-based models in which Ras adopts β1/2/3 orientations and binds to the Ras binding domain (RBD) of the kinase A-Raf (PDB:2MSE) (44). In these models, cationic side chains of K66, R68, and K69 on the A-Raf RBD are positioned to make favorable interactions with anionic membrane lipids. The potential binding of the B-Raf RBD to a similar β1/2/3 orientation of Ras has also been identified in molecular simulation (69). It is compelling that the modeled demarcation between membrane activity as a competitive inhibitor or a possible allosteric modulator straddles the β1/2/3 basin for many isolated RBDs (Figs. 2E, 2F, and 8). The potentially dramatic functional effects of subtle orientational modulation within the β1/2/3 basin are also apparent in Fig. 2C, where an 8-10° decrease in average tilt and a 6-14° decrease in average rotation convert “occluded” orientations that cannot bind the A-Raf RBD to “semi-exposed” orientations in which this binding is possible and perhaps even favorable (44, 48). The cell membrane’s ability to alternatively obstruct Raf binding to Ras or arrange both proteins for orientationally selected binding is illustrated in Fig. 9.

**Figure 9:**
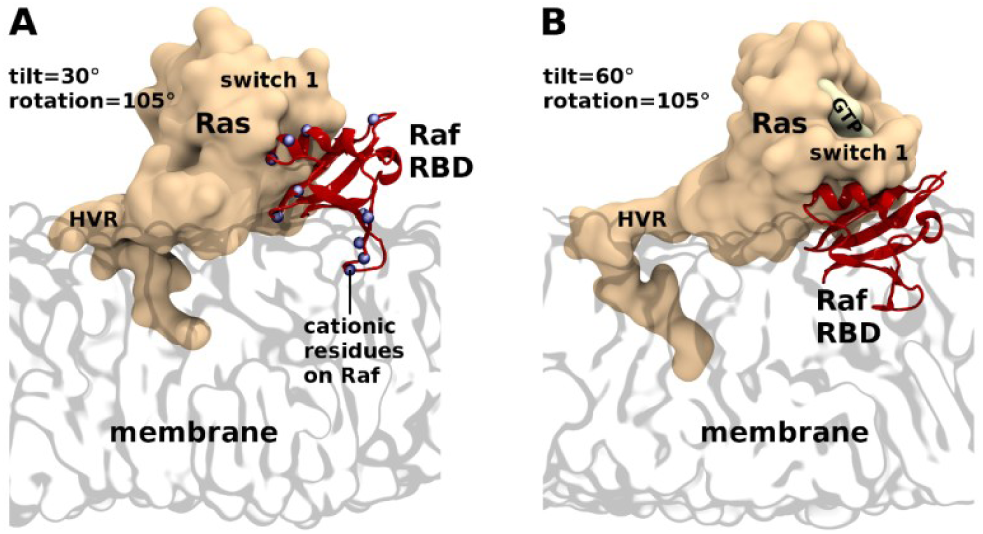
Modeled C-Raf RBD bound to simulated Ras in the β1/2/3 state. (A) Ras tilting by 30° permits Raf RBD binding with lipid head group immersion of RBD loop 4, which carries a positive charge in A- and C-RAF and a negative charge in B-RAF (90). (B) Ras tilting by 60° precludes RBD binding. RBD models mimic the 4G0N H-Ras/C-Raf complex (3).

## Conclusions

Extensive simulations reveal the impact of lipid composition and protein activation state on the behavior of full length K-Ras4B. Contact with anionic PS lipids draws Ras’ HVR and G domain toward the membrane surface and constricts the sampling of G domain orientations, consistent with previous simulations and data-driven models. In particular, PS lipids increase the population of orientations that bring beta sheets 1-3 and switch 1 toward the membrane surface. This orientational basin straddles a dividing line that separates dispositions of Ras that can bind to a variety of effector proteins from those in which binding is inhibited by the membrane itself. Therefore, existing compounds that stabilize this state (48) may be better able to block Ras signaling if they can be modified to promote slightly more tilting of the G domain toward switch 1. In addition to slowing Ras’ translational diffusion by a factor of two, PS lipids slow G domain reorientation by an order of magnitude and we show that the dominant pathway for G domain reorientation involves its transient escape from the membrane surface.

Conversely, this extensive set of GDP- and GTP-bound simulations does not reveal any statistically significant effects of the bound nucleotide on lateral or rotational diffusion, G domain orientation, or protein-lipid interactions. However, the computed half-life of active-state switch 1 configurations is up to two orders of magnitude shorter than observed in experiment, suggesting a problem with parameters or model form (potentially related to the magnesium ion), or that crystal structures identified as state 2 may not in fact represent Ras’ fully active configuration.

We also identify a rare event, stabilized by membrane embedding of switch 2, and correlating with C-terminal extension of G domain helix 5, that effectively hides Ras’ binding interfaces for many regulator and effector proteins. We suggest that this orientation represents another viable target for turning off Ras signaling by targeted stabilization of membrane adhesion.

## Author Contributions

CN and AEG designed research and wrote the manuscript. CN performed research and analyzed data.

## Acknowledgements

We thank Timothy Travers for providing CHARMM GDP and GTP parameters in GROMACS format, and Priyanka Prakash and Alemayehu A. Gorfe for providing structural models of their OS1 and OS2 states of K-Ras. This work has been supported in part by the JDACS4C program established by the U.S. Department of Energy (DOE) and the National Cancer Institute of the National Institutes of Health. This work was performed under the auspices of the U.S. DOE by Lawrence Livermore National Laboratory under Contract DE-AC52-07NA27344, Los Alamos National Laboratory (LANL) under Contract DE-AC5206NA25396, Oak Ridge National Laboratory under Contract DE-AC05-00OR22725, and Frederick National Laboratory for Cancer Research under Contract HHSN261200800001E. CN held a Director-funded postdoctoral fellowship from LANL. AEG was partially funded by U.S. DOE LDRD funds. Computations used resources provided by the LANL Institutional Computing Program, which is supported by the U.S. DOE National Nuclear Security Administration under Contract No. DE-AC52-06NA25396.

